# Supraphysiological Estradiol During Ovarian Stimulation Reveals Microbiome Resilience and *Prevotella* Opportunism

**DOI:** 10.64898/2026.06.02.729571

**Authors:** Maria J. Rus, Jack Lynch, Ana T. Marcos, Thuy Do, David Alarcón-Alarcón, José Manuel Navarro-Pando, Aurea Simon-Soro

## Abstract

How the oral ecosystem responds to acute, high-amplitude hormonal perturbations remains poorly defined. Here we exploit controlled ovarian hyperstimulation (COH), a pharmacologically defined model that elevates systemic estradiol (E2) ∼100-fold above physiological baseline to dissect the multi-layer response of the oral environment in ten healthy oocyte donors. Integrating paired serum and salivary hormone quantification with subgingival metatranscriptomics, we identify a dual principle governing the oral ecosystem under acute endocrine stress. First, salivary E2 reliably tracks intra-individual systemic dynamics, confirmed by robust linear regression, establishing saliva as a non-invasive endocrine proxy even under supraphysiological conditions. Second, despite the dramatic hormonal surge, the subgingival metatranscriptome exhibited functional resilience with no differentially expressed genes detected across two independent bioinformatic pipelines while a core of 20,687 genes remained stably expressed at both timepoints, underscoring the robustness of established biofilm communities. This global stability was selectively broken by four *Prevotella* species, whose abundance increased proportionally to salivary E2 elevation, consistent with their steroid hormone auxotrophy. Together, these findings reveal that the subgingival microbiome is functionally buffered against acute hormonal perturbation yet harbors estrogen-sensitive taxa capable of exploiting endocrine surges as ecological opportunities, with potential implications for reproductive and systemic inflammatory health.

## Introduction

Estradiol (E2) is a master regulator of female physiology, coordinating reproduction, immune modulation, vascular tone, and mucosal homeostasis across multiple organ systems(1, 2). Beyond its systemic roles, E2 exerts direct effects on peripheral mucosal environments, including the oral cavity, where clinical manifestations of hormonal sensitivity are well documented: gingival inflammation increases during pregnancy and the luteal phase,(3, 4) oral discomfort accompanies menopause, and cyclical shifts in subgingival microbial composition track the menstrual cycle(3, 5). These observations collectively position the oral ecosystem as a hormone-responsive interface, yet the biological mechanisms underlying its sensitivity to endocrine fluctuations remain incompletely defined.

The relationship between hormonal status and the oral microbiome has been explored primarily in the context of chronic or gradual transitions. Studies of menopause, hormone replacement therapy, and oral contraceptive use have demonstrated that sustained shifts in circulating estrogens reshape oral microbial communities, alter salivary composition, and modulate local immune tone(6). A recent synthesis of this evidence has framed the hormonal environment as a key determinant of oral microbiome structure across the female lifespan, identifying estrogen-sensitive taxa and niche-specific hormonal effects across saliva, tongue, and subgingival compartments(6). In the reproductive context, distinct gastrointestinal and reproductive microbial signatures have been identified in women undergoing assisted reproduction(7), suggesting that the hormonal milieu of the reproductive cycle shapes microbial communities beyond the gut and may contribute to infertility outcomes. However, these studies have largely examined compositional endpoints under chronic hormonal conditions. How the oral microbiome responds at the functional transcriptional level to acute, high-amplitude endocrine perturbations remains unknown.

This gap reflects in part a methodological challenge: acute hormonal events in humans are difficult to study prospectively because they are transient, variable in timing, and confounded by concurrent physiological changes. Controlled ovarian hyperstimulation (COH) offers a unique solution(8, 9). During COH, exogenous gonadotropins trigger rapid, supraphysiological E2 surges that exceed natural cycle peaks by several-fold, reaching concentrations associated with measurable local effects in oral tissues(3–5, 10, 11). This pharmacologically controlled trajectory provides a rare opportunity to study the oral ecosystem’s response to a quantified hormonal surge within a paired longitudinal design, under daily clinical monitoring and with a precisely defined temporal structure(12, 13). The oral cavity is particularly well suited for this investigation: saliva offers a non-invasive window into systemic endocrine dynamics through passive diffusion of free steroid hormones, while the subgingival niche, bathed by serum-derived gingival crevicular fluid and in direct contact with host vasculature, represents a microbial community with privileged exposure to circulating hormones.

Here, we hypothesized that salivary E2 would track systemic fluctuations while the subgingival microbiome would exhibit functional responses to the acute hormonal surge. To test this, we conducted a longitudinal study in women undergoing COH, integrating three complementary data layers: systemic and salivary E2 quantification to establish hormonal reflection across compartments, subgingival community profiling to assess compositional responses, and paired metatranscriptomics to resolve functional gene-level activity before and at peak stimulation. This multi-layer design allows us to distinguish between hormonal sensing at the biochemical level, ecological reorganization at the compositional level, and transcriptional reprogramming at the functional level. These three responses need not co-occur, and their dissociation is itself informative. We report that the oral ecosystem exhibits a dual principle under acute endocrine perturbation: global functional resilience at the metatranscriptomic level, coupled with selective compositional opportunism by estrogen-sensitive Prevotella taxa, with implications extending beyond oral health.

## Material and Methods

### Study Design and Cohort

This prospective longitudinal study was designed to assess whether a pharmacologically induced elevation in systemic estradiol levels could elicit detectable local changes in the oral environment. Saliva and subgingival plaque were selected as study matrices to represent two complementary levels of local reflection: saliva as a dynamic, hormone-sensitive fluid, and subgingival plaque as a stable microbial biofilm with close vascular and immune engagement. The study protocol received ethical approval from the Ethics Committee of the Universidad Europea del Atlántico (UNEATLANTICO) (Approval Number: CEI-24/2022). All procedures adhered to the principles of the Declaration of Helsinki, and written informed consent was obtained from all participants.

The study cohort included ten healthy female volunteers aged 20 to 35 years with regular menstrual cycles. All participants were undergoing COH solely for the purpose of oocyte donation, and gynaecologists at the collaborating fertility clinic confirmed the absence of underlying gynaecological disorders. Exclusion criteria included recent (within one month) use of antibiotics, antifungals, or oral antiseptics, and any current use of hormonal or systemic medications.

Each participant was followed over the course of one COH cycle. All individuals entered the study during the early follicular phase of their menstrual cycle (typically Day 2 or 3). Baseline evaluations (designated T1) were performed prior to hormonal stimulation and included both systemic and oral assessments. Systemic measurements at T1 comprised serum levels of Follicle-Stimulating Hormone (FSH), Luteinizing Hormone (LH), and Estradiol (E2). Oral assessments at T1 included a comprehensive periodontal evaluation and the collection of unstimulated saliva and subgingival plaque, with the respective procedures detailed below in *Sample Collection* and *Clinical Data Collection*.

Hormonal stimulation was initiated after baseline sampling and included daily subcutaneous administration of gonadotropins: Gonal-f®, a recombinant FSH, and HMG Lepori®, containing both FSH and LH activity. The initial dosage was determined based on baseline ovarian reserve parameters (FSH, LH, E2 levels). Dosage was then adjusted dynamically throughout the stimulation phase. These adjustments were guided by daily monitoring of serum estradiol concentrations and assessment of follicular development via transvaginal ultrasonography, which included measurement of endometrial thickness. Consequently, while the types of hormones used were consistent, the total cumulative dose and the exact duration of stimulation varied among participants based on individual response.

A second set of oral samples was obtained when oocytes were deemed mature for transvaginal retrieval, designated T2. This T2 collection point also corresponded to the peak systemic estradiol response and occurred approximately 13 days after T1. The same oral sampling procedures used at baseline were employed at T2.

### Sample Collection

Oral samples were collected from each participant at two time points: T1 (baseline, prior to hormonal stimulation) and T2 (day of oocyte retrieval). At both visits, unstimulated saliva and subgingival plaque were obtained.

For saliva collection, participants were instructed to refrain from eating, drinking, or performing oral hygiene procedures for at least one hour prior to sampling. They were then asked to passively drool into a sterile 50 mL collection tube over a continuous 5-minute period.

Subgingival plaque was collected from all teeth present using a sterile Gracey curette applied in a disto-mesial sweep along both buccal and lingual surfaces. The collected plaque material from all sites was pooled into a single cryotube containing 1 mL of RNA Shield™ (Zymo Research, Irvine, CA, USA), a stabilizing reagent that preserves microbial RNA. All samples were immediately placed on ice after collection and stored at −80°C within two hours to ensure preservation of RNA integrity for subsequent metatranscriptomic analysis.

Peripheral blood samples were obtained for hormonal analysis as part of routine clinical monitoring during the COH protocol. Serum levels of FSH were measured at T1 (baseline). LH concentrations were assessed at T1 and on the day prior to oocyte retrieval. E2 levels were measured at T1 and subsequently monitored approximately daily throughout the stimulation phase until the day before oocyte retrieval, guiding gonadotropin dose adjustments and the trigger for final oocyte maturation.

### Clinical Data Collection

General systemic health information was obtained from each potential participant using a guided health questionnaire. This included self-reported history of systemic diseases, current medication use (hormonal and non-hormonal), and lifestyle factors (alcohol consumption, tobacco use, other drug use). This screening confirmed eligibility based on the study inclusion/exclusion criteria before participants were formally enrolled and proceeded to baseline assessments.

Oral clinical data included a periodontal evaluation performed at the baseline visit (T1) using the Basic Periodontal Examination (BPE) protocol (14, 15). Periodontal status was characterized not only to assess baseline oral health but also to contextualize potential microbial shifts within any pre-existing inflammatory environment, which could modulate the local response to systemic hormonal changes.

For the BPE assessment, the dentition was divided into canine-limited sextants, excluding third molars. A sextant containing only one tooth was excluded, and its tooth was evaluated as part of an adjacent sextant. A calibrated, mm-marked periodontal probe was introduced between the tooth and the gingiva to determine the depth of the gingival sulcus or pocket relative to the level of the gingival margin. Probing was performed at six points on each tooth: mesial, midpoint, and distal of the vestibular and palatal/lingual side. Bleeding on probing (BoP), the presence of supra- and subgingival calculus, plaque-retentive factors (e.g., restoration overhangs), furcation involvement, and clinical attachment loss (CAL) were also recorded. The BPE score for each sextant was determined by the highest probing depth or the most severe clinical finding recorded for any tooth within that sextant.

Based on the highest BPE scores across the sextants, participants were categorized according to their overall periodontal status, which reflected diagnosis and treatment needs. A *healthy periodontium* was defined when all sextant scores indicated the absence of BoP, calculus, and defective margins, with probing depths remaining ≤3.5 mm. *Gingivitis* was characterized by the presence of BoP, calculus, or plaque-retentive factors, while probing depths were still ≤3.5mm. Participants were classified with *moderate periodontitis* if the highest scores indicated probing depths between 4 mm and 5.5 mm in one or more sextants. Finally, *severe periodontitis* was assigned if the highest score revealed probing depths ≥6 mm in at least one sextant, or if furcation involvement or CAL ≥7 mm was present. This baseline periodontal classification characterized the initial oral health status of the cohort and served as a key clinical variable in subsequent analyses.

Relevant gynaecological data were also collected, including history of previous pregnancies and prior oocyte donation cycles. As part of standard clinical monitoring during COH, endometrial thickness was measured via transvaginal ultrasonography at baseline (Day 0) and approximately every two days thereafter until the day before oocyte retrieval.

Clinical outcome data related to the donation cycle and recipient outcomes were recorded. These included: total number of oocytes retrieved, number of mature (MII) oocytes, number fertilized, number of embryos transferred to the recipient, number of viable embryos cryopreserved, and the resulting pregnancy outcome in the recipient (if applicable/available).

### Salivary Estradiol Measurement

The concentration of 17β-estradiol was measured in unstimulated saliva samples collected at baseline (T1) and on the day of oocyte retrieval (T2). Measurements were performed using a commercial enzyme immunoassay (ELISA) kit validated for salivary analysis (Salimetrics®, State College, PA, USA), following the manufacturer’s protocol. Samples were assayed in duplicate. Final concentrations were determined by averaging the duplicate optical density readings and interpolating the result against a standard curve generated during the same assay run. Salivary estradiol values were reported in picograms per milliliter (pg/mL).

### RNA Extraction and Metatranscriptomic Sequencing

Total RNA was extracted from subgingival plaque samples preserved in RNA Shield™ using the RNeasy® PowerMicrobiome® Kit (QIAGEN, Hilden, Germany). The extraction followed the manufacturer’s protocol, optimized for microbial RNA recovery from complex biological samples. An on-column DNase I treatment step was included to eliminate contaminating genomic DNA. RNA quantity was determined using a Qubit™ Fluorometer (Thermo Fisher Scientific, Waltham, MA, USA), and RNA integrity was assessed using an Agilent 2100 Bioanalyzer system (Agilent Technologies, Santa Clara, CA, USA).

Ribosomal RNA (rRNA) depletion, library preparation, and sequencing were outsourced to Macrogen Inc. (Seoul, Republic of Korea). Briefly, bacterial rRNA was depleted from the extracted total RNA using the NEBNext® rRNA Depletion Kit (Bacteria) (New England Biolabs, Ipswich, MA, USA). Subsequently, stranded mRNA sequencing libraries were constructed using the TruSeq® Stranded Total RNA Library Prep Gold Kit (Illumina, Inc., San Diego, CA, USA), following the manufacturer’s guidelines (Reference Guide 1000000040499 v00). The final libraries were sequenced on an Illumina NovaSeq 6000 platform, generating 151 bp paired end reads.

### Bioinformatic Analysis of Metatranscriptomes

The bioinformatic analysis of the raw metatranscriptomic sequencing data involved a sequential pipeline encompassing quality control (QC), host and rRNA filtering, mapping, quantification, differential expression (DE) analysis, and functional annotation. Raw paired-end reads (FASTQ format) underwent initial QC assessment using FastQC (v0.12.1) (16). Reads were then processed with fastp (v0.23.4) (17) for adapter trimming, quality filtering (Phred score ≥ 20) and removal of reads shorter than 70 bp. To remove human host reads, filtered reads were aligned to the human reference genome (Gencode Release 47, GRCh38.p14) using Bowtie2 (v2.5.4) (18) with the --very-sensitive preset. Unmapped read pairs were retained. Residual rRNA reads were identified and removed from the non-host reads using SortMeRNA (v4.3.7) (19) against its default bacterial rRNA databases (SILVA, Rfam). Reads not matching rRNA were kept for downstream analysis.

Cleaned, non-rRNA reads were mapped to the Human Oral Microbiome Database (HOMD, RefSeq v10.1) (20), annotated using PROKKA (v1.14.6) (21). Given that HOMD predominantly comprises bacterial and, to a lesser extent, archaeal genomes, the subsequent transcriptional analysis primarily reflects the activity of these prokaryotic domains, with a principal focus on the bacterial component. Mapping was performed using Bowtie2 (v2.5.4) in paired-end mode with the --very-sensitive preset, generating BAM alignment files. Gene expression levels were quantified from BAM files using featureCounts (subread package v2.0.6) (22). Reads were assigned to coding sequences (CDS) based on the HOMD PROKKA GTF annotation file (-t CDS), summarized per gene locus tag (-g locus_tag). Paired-end reads were counted (-p --countReadPairs) and reads mapping to multiple features were fractionally counted (-M). This produced a gene-by-sample count matrix, which was then used for differential gene expression analysis as detailed below in *Statistical Analysis*.

### Validation via Alternative Pipeline

To validate the primary findings and broaden the taxonomic scope of the analysis, a secondary, independent pipeline was employed using a custom-built protein database. Using similar methodologies as previously described for the detection of oral archaea in oral DNA and RNA samples, this complementary approach was designed to overcome the prokaryote-centric limitation of the HOMD-based analysis by explicitly including viral, archaeal and eukaryotic organisms previously reported to colonise the human oral cavity (23).

In brief, a bespoke protein database was constructed using DIAMOND (makedb function). This database integrates protein sequences from several sources, including all microbes from the HOMD, as well as archaea, viruses, and eukaryotes previously identified in the human oral cavity, as previously reported (23–26). The viral component of the database included both human viruses and bacteriophages to ensure a complete view of the oral virome (27, 28).

For this validation analysis, the raw sequencing reads were processed through a distinct filtering workflow. Initial quality was assessed with FastQC (0.12.0), after which reads were processed with cutadapt (v5.0). The trimming parameters involved removing 10 base pairs from both the 5’ and 3’ ends of each read, followed by the removal of any read shorter than 30 base pairs. These processed reads then underwent host removal by alignment to the human genome with Bowtie2, as previously, and the resulting unmapped reads were used for microbial analysis. The non-host reads were mapped against the bespoke database using DIAMOND (v2.0.15). To ensure high confidence in alignments assignments, a stringent filter was applied, retaining only alignments with ≥97% sequence identity and ≥90% query coverage. A gene-by-sample count matrix was generated from these filtered hits and Counts Per Million mapped reads (CPM) was calculated for each alignment. Subsequently, count matrices were imported into R for analysis with packages such as DESeq2, as described below. This complementary method provides a more holistic view of the active oral microbiome and serves as a crucial validation for the results obtained from the HOMD-based workflow.

### Statistical Analysis

All statistical analyses were performed using R (v4.4.1) (29). Descriptive statistics (median, interquartile range [IQR] or mean ± standard deviation [SD], as appropriate) were calculated for baseline clinical and hormonal variables. Changes in serum and salivary estradiol levels between T1 and T2 were assessed using the paired Wilcoxon signed-rank test, chosen due to the small sample size and potential non-normality of hormonal data distribution.

Correlations between continuous variables (salivary vs. serum estradiol at T1 and T2) were evaluated using Spearman’s rank correlation coefficient (ρ). A correlation matrix visualizing pairwise associations among key participant characteristics, treatment variables, and estradiol levels (e.g., Age, Periodontal Status, total gonadotropin doses, oocyte yield, and serum/salivary E2 levels at T1 and T2) was generated using Spearman coefficients.

To investigate the relationship between the change in serum estradiol (ΔSerum E2 = T2 - T1) and the corresponding change in salivary estradiol (ΔSaliva E2 = T2 - T1), a robust linear regression model was fitted using the rlm() function from the MASS package (30). This model adjusted for potential confounders: total cumulative doses of Gonal-f® and HMG Lepori®, and the duration of stimulation (days). Robust regression was selected to minimize the influence of potential outliers in this small cohort. Model fit was assessed via standard residual diagnostic plots. The regression coefficient (β) and associated *p*-value for the primary predictor (ΔSerum E2) were reported. All statistical tests were two-tailed, and a *p*-value < 0.05 was considered statistically significant. Figures were generated using the R package ggplot2 (31).

Furthermore, a correlation analysis was performed to investigate the relationship between clinical parameters and the abundances of dominant microbial strains at three distinct levels:

(1) *Baseline*, correlating T1 clinical variables with T1 microbial abundances; (2) *Oocyte retrieval*, correlating T2 clinical variables with T2 microbial abundances; and (3) *Change (*Δ*T2-T1)*, correlating the change in clinical variables with the corresponding change in microbial strain abundances. For this analysis, the top 15 most abundant microbial strains, determined by their mean abundance across all samples, were selected. Spearman’s rank correlation coefficients (ρ) and corresponding p-values were calculated between these 15 strains and the relevant clinical variables for each of the three levels using the Hmisc (32) package in R. The resulting correlation matrices were visualized as a heatmap using the pheatmap package (33). To highlight significant relationships, individual cells in the heatmap corresponding to a p-value < 0.05 were annotated with an asterisk (*).

Differential gene expression (DGE) analysis of the subgingival metatranscriptomes, comparing T2 against T1, was conducted using the DESeq2 package (v1.44.0) (34). Prior to DGE analysis, genes with low counts (defined as those with fewer than 10 counts in at least 5 samples) were filtered out. Subsequently, Principal Component Analysis (PCA) was performed on Variance-Stabilized Transformed (VST) count data. This was used to visualize the global structure of the metatranscriptomic data, assess the primary sources of variance among samples, and confirm the appropriateness of a paired statistical design by examining sample clustering (e.g., by subject, timepoint, periodontal status). For the DGE analysis, a design formula of ∼SubjectID + Timepoint was used to account for the paired nature of the samples and inter-individual variability. The DESeq2 pipeline involved normalization of gene counts using the median-of-ratios method, estimation of gene-wise dispersions, and fitting a negative binomial generalized linear model. Statistical significance of differential expression was assessed using the Wald test. Results were extracted for the contrast between T2 and T1, with T1 as the reference level. Genes with a Benjamini-Hochberg adjusted *p*-value (FDR) < 0.05 were considered significantly differentially expressed (35). Volcano plots were generated using the EnhancedVolcano package (36) to visualize DGE results.

### Data and Software Availability

The raw RNA sequences and associated metatranscriptomic metadata have been deposited in the European Nucleotide Archive (ENA) under accession number PRJEB101296. The R code and scripts utilized for the pre-processing, statistical analyses, and figure generation are publicly accessible via Zenodo, DOI: 10.5281/zenodo.17416734.

## Results

### Acute Systemic Estradiol Surge During COH is Reflected Locally in Saliva

To investigate whether the pronounced systemic hormonal shifts induced by Controlled Ovarian Hyperstimulation (COH) manifest locally in the oral environment, we first tracked estradiol concentrations in both serum and saliva (see study workflow in Figure 1A). As anticipated from the COH protocol, serum estradiol levels underwent a dramatic and statistically significant increase from baseline (T1, early follicular phase) to the day of oocyte retrieval (T2) (Figure 1B, Left panel; paired Wilcoxon test, *p* = 0.006). Crucially, this systemic surge was mirrored locally, with salivary estradiol concentrations also showing a significant elevation between T1 and T2 (Figure 1B, Right panel; paired Wilcoxon test, *p* = 0.008), providing initial evidence for systemic-to-oral hormonal reflection.

**Figure 1.**
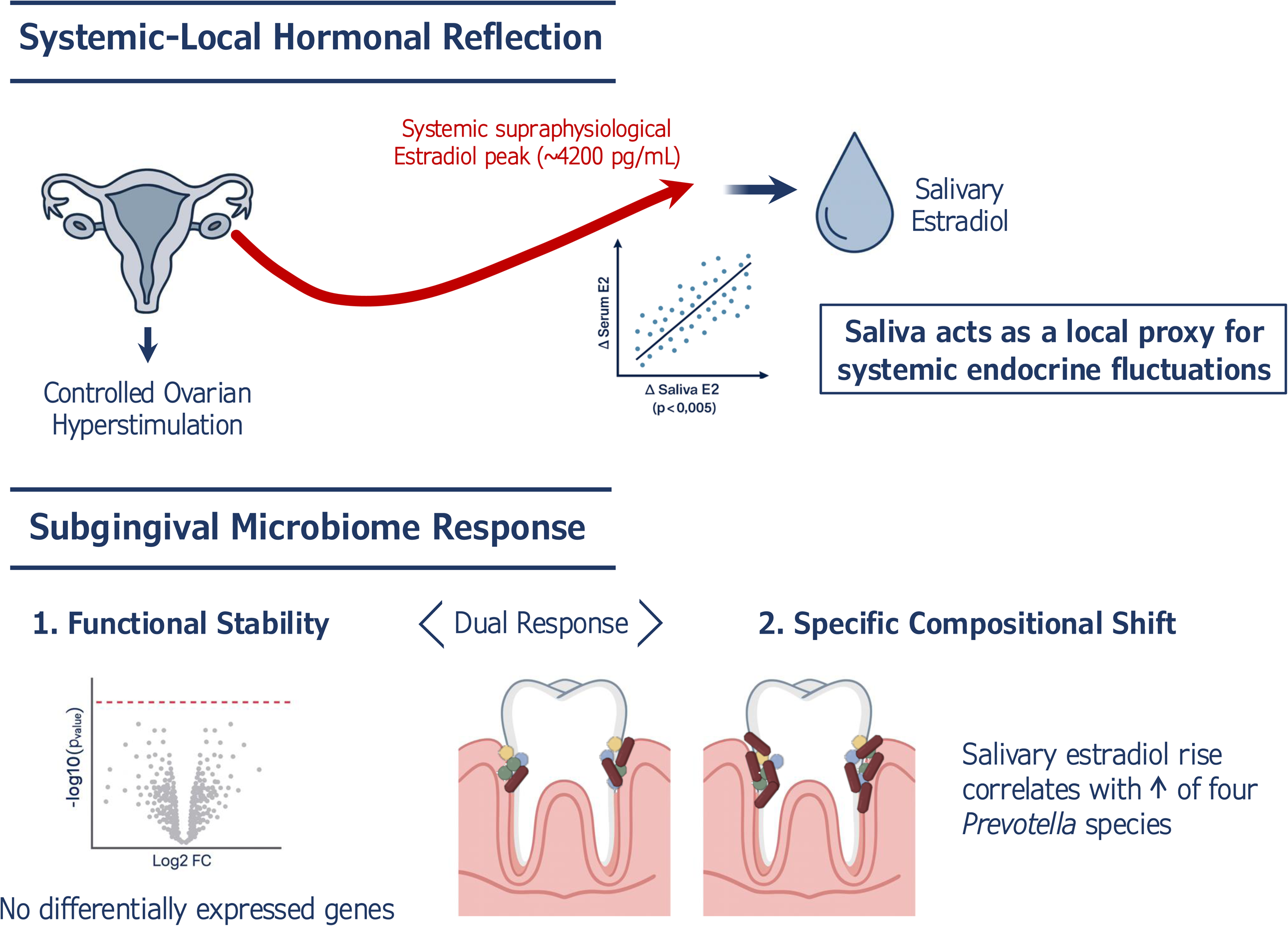
Systemic Estradiol (E2) Surge During COH is Dynamically Reflected in Saliva and Correlates with Clinical Parameters. a) Experimental design for assessing oral responses to COH-induced systemic E2 elevation in 10 healthy female oocyte donors. The diagram outlines the study progression, detailing sample collection and key monitored parameters at two primary timepoints: T1 (baseline, prior to hormonal intervention) and T2 (day of oocyte retrieval, corresponding to peak E2 response and oocyte maturation). At T1 (Baseline): Systemic data collection encompassed initial serum levels of E2 (∼40 pg/mL, reflecting baseline physiological values of the early follicular phase). Oral samples included saliva (for estradiol measurements) and subgingival plaque (for metatranscriptomic analysis). COH Protocol: Following T1 assessments, participants underwent daily gonadotropin stimulation. Throughout this phase, systemic E2 levels and endometrial thickness were monitored to guide adjustments to the stimulation protocol and determine the optimal timing for inducing final oocyte maturation. At T2 (Oocyte Retrieval): Systemic data collection was repeated for final E2 serum levels (reaching peak levels of ∼4500 pg/mL, far exceeding the normal physiological range). Oral samples were collected again for the same analyses as T1. b) Comparison of E2 levels in serum (left) and saliva (right) between baseline (T1, early follicular phase) and the day of oocyte retrieval (T2). Boxplots depict the median and interquartile range (IQR), with individual data points overlaid for each participant (circles for T1, squares for T2; n=10). Statistical significance was determined using a paired Wilcoxon signed-rank test, with exact p-values shown above the comparison bars. c) Correlation between serum and saliva E2 levels at baseline T1 (left) and on the day of oocyte retrieval T2 (right). Each point represents an individual participant. Spearman’s correlation coefficient (ρ) and the corresponding p-value are displayed in the upper corner of each plot. d) Relationship between the magnitude of change in serum E2 levels (Δ Serum E2 = T2 - T1) and the corresponding change in saliva (Δ Saliva E2 = T2 - T1). Each triangle represents an individual participant (n=10). The solid line indicates the robust linear regression fit, with the shaded area representing the 95% confidence interval. The regression coefficient (β) and t-statistic from the robust linear regression analysis are displayed. e) Heatmap visualizing Spearman’s rank correlation coefficients (ρ) between various participant characteristics and treatment-related variables (rows) and E2 levels in serum and saliva at T1 and T2 (columns). Color intensity and hue indicate the strength and direction of the correlation, according to the provided scale. Significant correlations *(p* < 0.05) are shown in bold.

We next explored the direct relationship between systemic (serum) and local (salivary) estradiol levels at each time point. Under baseline physiological conditions (T1), a significant positive correlation was observed (Figure 1C, Left panel; Spearman’s ρ = 0.72, *p* = 0.018), indicating that circulating estradiol levels are proportionally reflected in saliva under normal hormonal ranges. However, this relationship appeared to break down under the supraphysiological conditions at T2. The simple correlation between absolute serum and salivary estradiol levels was no longer statistically significant (Figure 1C, Right panel; Spearman’s ρ = 0.38, *p* = 0.28). Visual inspection revealed that while the highest serum levels corresponded to high salivary levels, participants with mid-range supraphysiological serum E2 (approx. 2000–3000 pg/mL) exhibited salivary concentrations compressed within a relatively narrow upper range (approx. 2.4–3.8 pg/mL). This attenuation in the dynamic scaling of salivary E2 may reflect compression effects, potentially due to physiological constraints or technical limitations at high systemic concentrations.

Participants exhibited considerable inter-individual variability in both their COH treatment parameters and hormonal responses, as summarized in Table 1. Gonadotropin exposure was heterogeneous across the cohort, both in terms of total administered doses (mean ± SD: Gonal-f® 1709.8 ± 874.05 IU; HMG Lepori® 1331.25 ± 220.26 IU) and stimulation duration (12.8 ± 1.87 days). This inter-individual variability was also evident in E2 levels. At T1, serum E2 concentrations showed considerably higher dispersion (serum 42.31 ± 21.16 pg/mL) than salivary levels (saliva 1.09 ± 1.22 pg/mL). This trend persisted at T2, where final serum estradiol levels varied widely among participants (4208.9 ± 3649.61 pg/mL), while salivary concentrations remained more tightly clustered (3.80 ± 1.95 pg/mL).

**Table 1.**
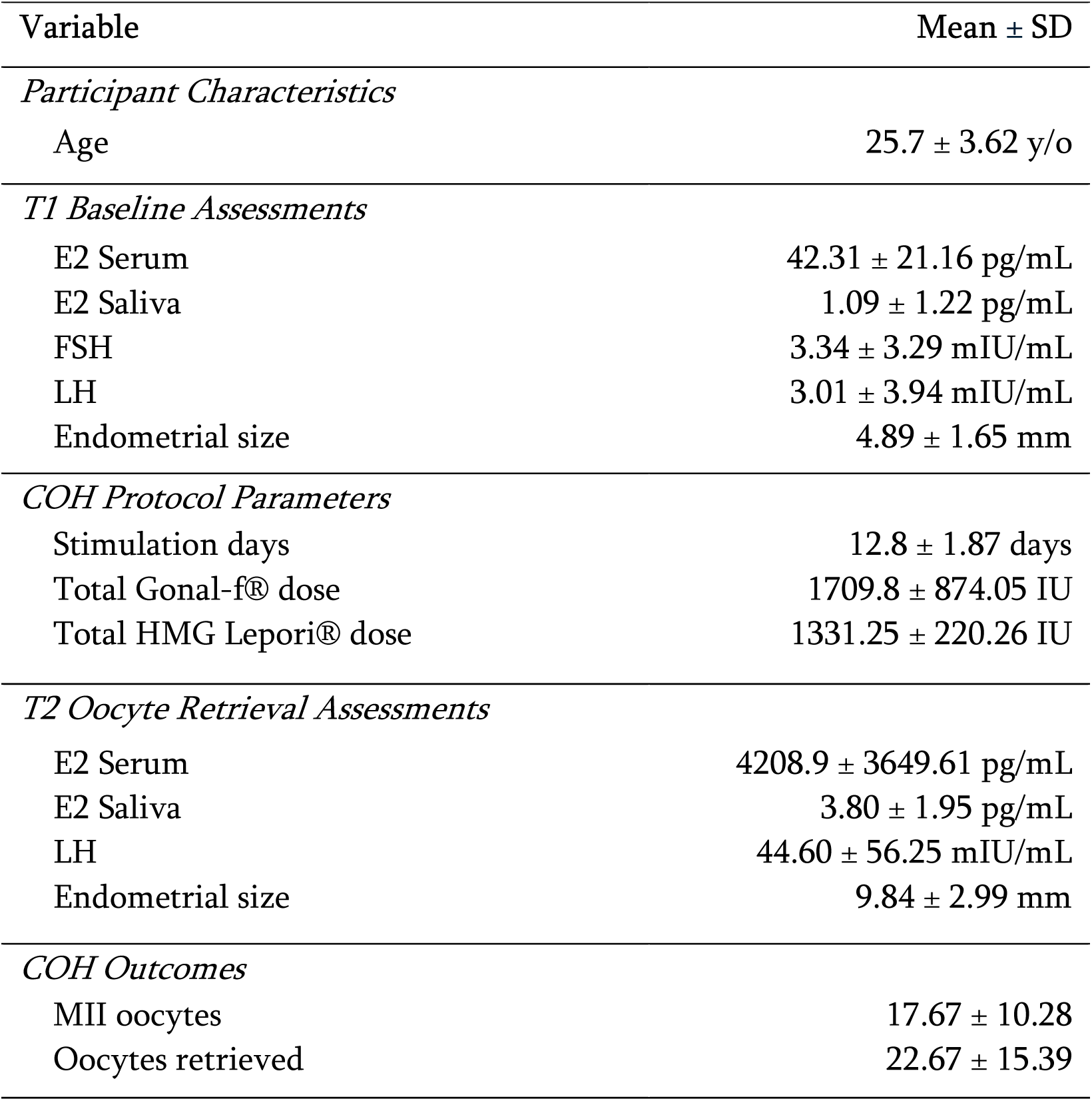
Descriptive statistics of study variables. The table presents mean ± SD for key variables collected during the study. This includes participant age, hormonal levels (E2, LH, FSH) in serum and saliva at T1 (baseline) and at T2 (oocyte retrieval), endometrial size, details of the stimulation protocol (stimulation days, Gonal-f® and HMG-Lepori® dose), and oocyte outcomes (total MII oocytes, total oocytes retrieved).

Given this inherent variability and the previously noted lack of simple correlation for absolute E2 levels at T2, we employed robust linear regression to determine if the magnitude of change in systemic estradiol (ΔSerum E2 = T2 - T1) was consistently associated with the corresponding change in salivary estradiol (ΔSaliva E2 = T2 - T1), while statistically controlling for the confounding effects of gonadotropin dosage and stimulation duration. This analysis revealed a significant positive association (Figure 1D; β = 0.0006, *t* = 4.16, *p* < 0.005). This finding strongly supports that, despite the complexities at peak levels, the dynamic systemic hormonal fluctuations induced by COH are indeed reliably reflected in the salivary compartment when treatment variations are accounted for.

Further analysis explored factors associated with the magnitude of the systemic hormonal response and potential confounders influencing local salivary levels. Consistent with the clinical role of estradiol as an indicator of ovarian activity during COH, peak serum estradiol levels at T2 demonstrated a strong positive correlation with the total number of oocytes retrieved and mature (MII) oocytes (Figure 1E; Spearman’s ρ = 1.00, *p* < 1.54 × 10⁻⁸ for both). In contrast, an inverse correlation was observed between the total administered dose of Gonal-f® and final serum estradiol concentrations (T2), where participants receiving higher cumulative doses tended to exhibit lower peak E2 levels (Figure 1E; Spearman’s ρ = −0.63, *p* = 0.05). This finding may reflect clinical dose adjustments based on anticipated ovarian response, highlighting the individualized nature of systemic reactions to stimulation.

Finally, to ascertain whether local factors influenced salivary estradiol concentrations under these high-estrogen conditions, we examined potential associations with participant characteristics. Exploratory correlation analyses revealed no statistically significant associations between salivary estradiol levels at T2 and either participant age (Spearman’s ρ = 0.03, *p* = 0.92) or baseline periodontal status as categorized by BPE scores (Spearman’s ρ = - 0.50, *p* = 0.44). The absence of these correlations suggests that, within this cohort, the measured increase in salivary estradiol predominantly reflected the systemic endocrine surge rather than being substantially modulated by participant age or pre-existing periodontal condition.

### Subgingival Microbial Community Shows Compositional Stability but Taxon-Specific Responses

Having established that the acute systemic estradiol surge is mirrored locally in saliva, we next investigated its impact on the composition of the subgingival microbial community. Community-level investigation using the customized multi-domain database supported the observation of overall stability. Alpha diversity metrics (Observed, Chao1, Shannon, and Simpson) showed no statistically significant differences between T1 and T2 in paired tests, indicating that the richness and evenness of the active microbial community remained unchanged (Figure 2A). Similarly, beta diversity, visualized with a Principal Coordinates Analysis (PCoA) of Bray-Curtis dissimilarity values, showed no distinct clustering by time point, with a PERMANOVA test confirming the lack of a significant community wide compositional change (p = 0.924) (Figure 2B).

**Figure 2.**
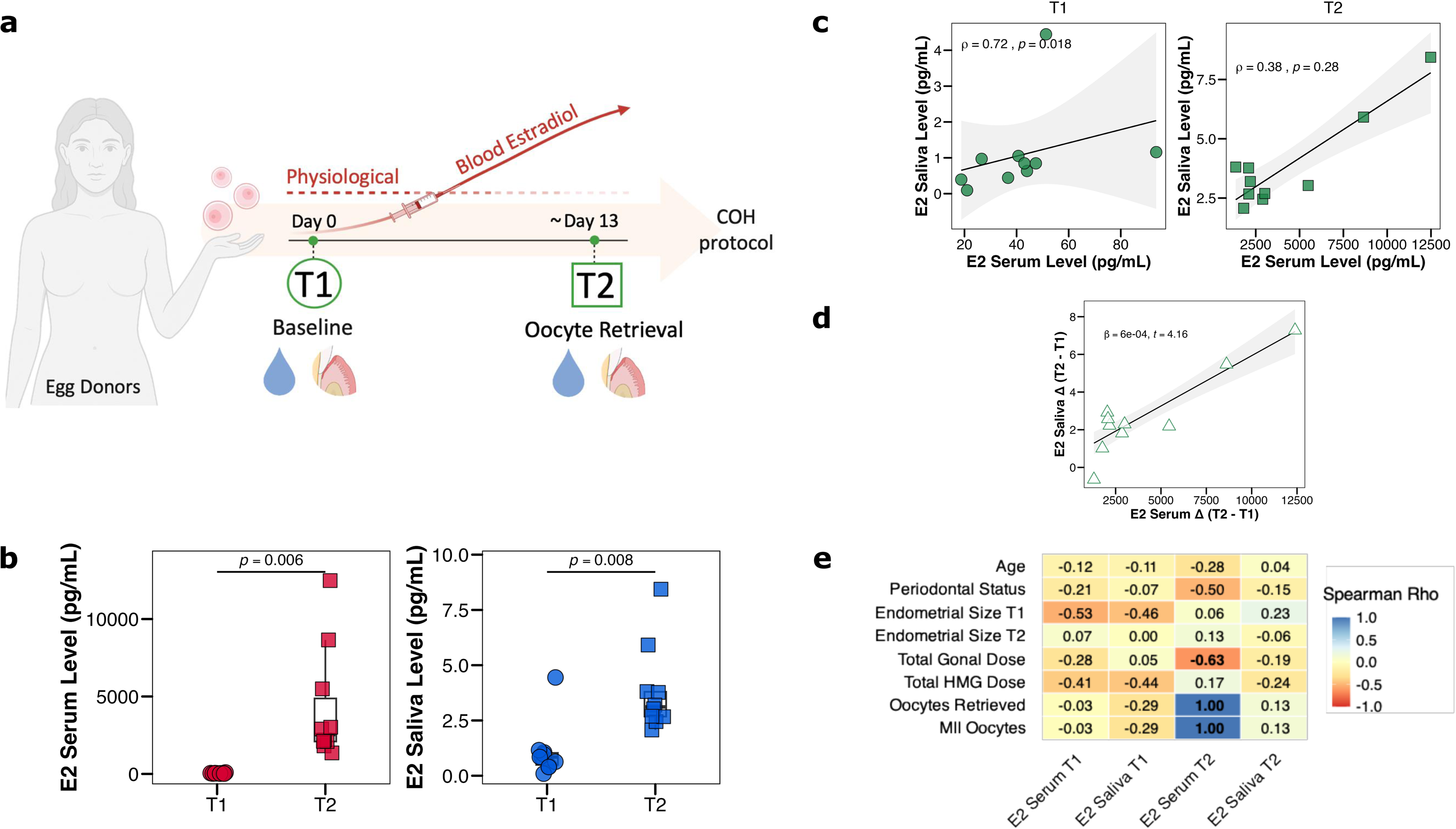
Stability of Subgingival Microbial Species Diversity Following Acute Estradiol Surge. a) Table summarizing the community composition at the domain level, showing the abundance of bacteria, eukaryota, viruses, and archaea identified from the custom oral database dataset. Values represent the mean relative abundance (%) ± Standard Deviation for all samples at T1 (Baseline) and T2 (Oocyte Retrieval). No significant differences were observed in any domain between timepoints, indicating a stable community structure at this high taxonomic rank (*p* > 0.05). b) Longitudinal analysis of intraindividual species diversity (alpha diversity) between T1 and T2. The plots display four standard metrics: Observed species and Chao1 for richness, and Shannon and Simpson for evenness. Each point represents a sample, and lines connect paired samples from the same participant. A Wilcoxon signed-rank test revealed no significant changes in any of the diversity indices (*p* > 0.05), indicating stability in the within-sample community structure. c) Principal Coordinate Analysis (PCoA) plot based on the Bray-Curtis dissimilarity matrix of microbial species counts, illustrating the community-level compositional differences (beta diversity) between timepoints. The plot shows the first two principal coordinate axes, with the percentage of total variance explained by each axis indicated (PCo1: 55.3%, PCo2: 11.9%). The overlapping 95% confidence ellipses and the lack of distinct clustering demonstrate high similarity in community structure between the two timepoints. This observation is statistically supported by a Permutational Multivariate Analysis of Variance (PERMANOVA), which found no significant difference in the overall community composition (*p* = 0.924). d) Heatmaps showing Spearman’s rank correlation coefficients (ρ) between selected clinical variables and the normalized abundances of the 15 most abundant microbial strains. The color scale indicates the strength and direction of the correlation (red, positive; blue, negative), with statistically significant relationships (p < 0.05) marked by an asterisk (*). The panel presents three distinct analyses: at baseline (T1), correlating clinical variables (Age, Periodontal Status, E2 Serum T1, E2 Saliva T1) with microbial abundances, where no significant correlations were found; at oocyte retrieval (T2), correlating clinical variables (Gonal Dose, HMG Dose, E2 Serum T2, E2 Saliva T2) with T2 abundances, which revealed a significant negative correlation between Gonal dose and *Streptococcus cristatus* AS 1.3089, and a positive correlation between salivary E2 and *Actinomyces oris*; and longitudinal changes (Δ T2–T1), correlating the change in serum and saliva estradiol with the corresponding change in strain abundance. This final analysis identified significant positive correlations between the change in salivary E2 and several *Prevotella* species, including *P. fusca* JCM 17724 and *P. denticola* F0289.

While the overall community composition remained stable, we performed an exploratory analysis to determine if the abundances of specific dominant microbial strains were associated with the clinical and hormonal parameters of the COH cycle. At baseline (T1), we found no significant correlations between clinical variables, including age, periodontal status, and E2 levels, and the abundances of the top 15 most abundant strains (Figure 2D). However, following the hormonal surge at T2, several significant associations emerged. Notably, the total administered Gonal-f® dose showed a significant negative correlation with the abundance of *Streptococcus cristatus* AS 1.3089, while salivary E2 levels were positively correlated with the abundance of *Actinomyces oris*. Furthermore, the analysis of longitudinal changes revealed that the magnitude of the increase in salivary E2 between T1 and T2 was significantly and positively correlated with a corresponding increase in the abundance of four *Prevotella* species: *P. melaninogenica* ATCC 25845, *P. scopos* JCM 17725, *P. fusca* JCM 17724, and the periodontitis-associated *P. denticola* F0289 (Figure 2D). These findings suggest that while the overall community function is resilient to acute hormonal shifts, a differential estrogen sensitivity may exist among specific bacterial taxa. In particular, certain *Prevotella* species appear responsive to the elevated local endocrine environment, exhibiting compositional changes not observed in other dominant community members.

### The Subgingival Metatranscriptome Demonstrates Functional Resilience Following Acute Estradiol Surge

Finally, to assess the functional consequences of the pronounced hormonal fluctuation, we investigated the transcriptional activity of the microbial community within the subgingival plaque. This niche, intimately exposed to gingival vasculature, represents a key interface for potential host-microbiome endocrine interactions.

To characterize the overall structure of microbial gene expression profiles and assess the impact of COH, we first performed Principal Component Analysis (PCA) on Variance-Stabilized Transformed (VST) counts of filtered genes. The PCA revealed that the dominant source of variation in the subgingival metatranscriptome was interindividual heterogeneity (Figure 3A). Samples clearly clustered by participant (indicated by color), demonstrating strong, individualized microbial transcriptional signatures that remained relatively stable across the study period. While intraindividual changes occurred, there was no consistent separation or trajectory distinguishing T1 samples from T2 samples across the cohort. This prominent intersubject variability highlights the personalized nature of the subgingival microbiome’s functional state and underscores the necessity of using paired statistical models to control for baseline differences when assessing intervention effects.

**Figure 3.**
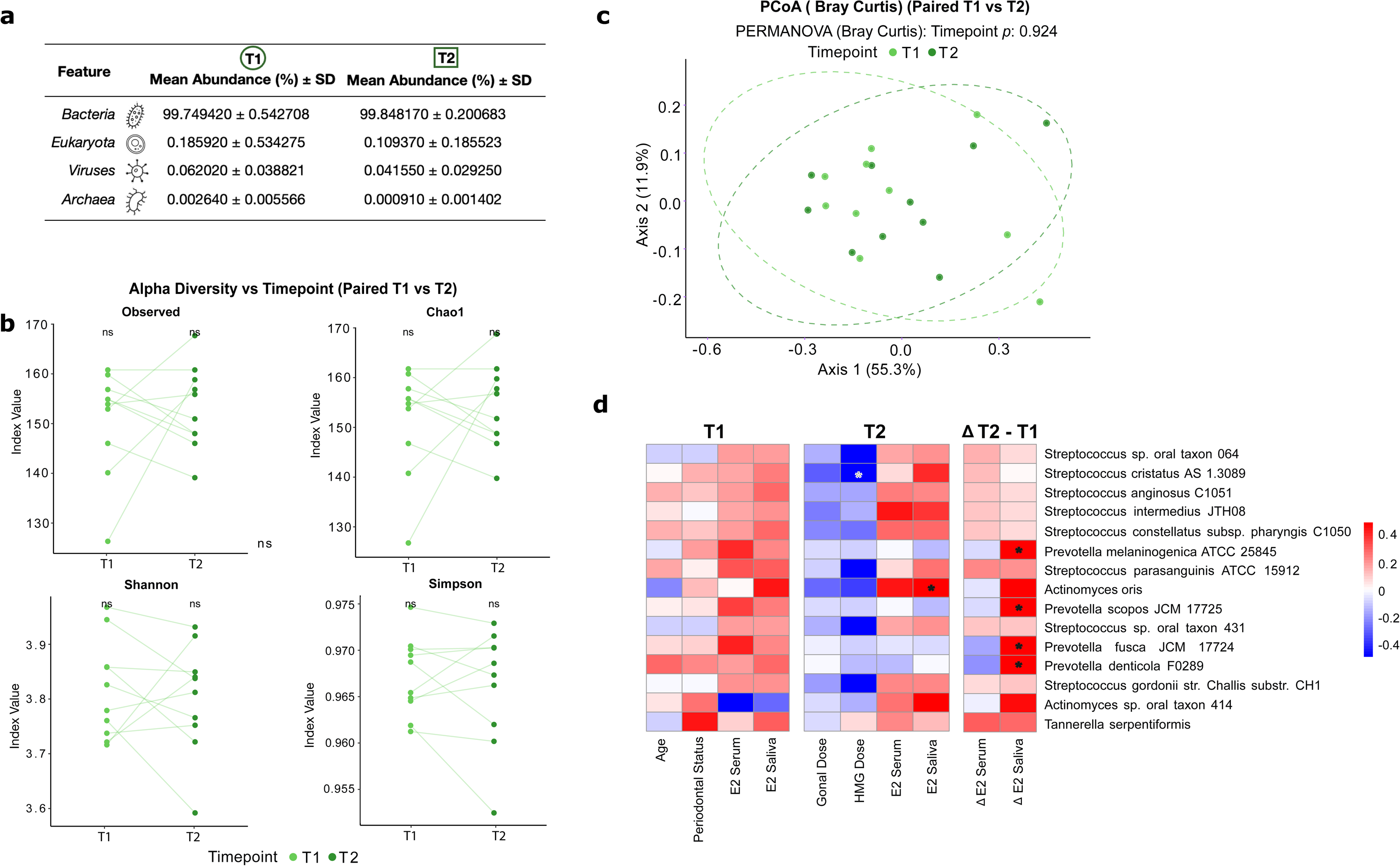
Functional Resilience of the Subgingival Microbial Community in Response to Acute Estradiol Elevation. a) Principal Component Analysis (PCA) plot based on Variance-Stabilized Transformed (VST) counts of filtered genes from the subgingival metatranscriptome. The plot shows the first two principal components, with the percentage of total variance explained by each axis indicated (PC1: 49%, PC2: 12%). Each point represents a sample, colored according to Subject ID and shaped according to the timepoint (circles: T1, baseline; triangles: T2, day of oocyte retrieval). The visualization highlights strong clustering by participant, indicating pronounced interindividual heterogeneity and intraindividual stability in microbial gene expression profiles. b) Volcano plot displaying the results from the DESeq2 paired differential expression analysis of subgingival metatranscriptomic data, comparing the day of oocyte retrieval (T2) versus baseline (T1). The x-axis represents the Log2 Fold Change in gene expression, and the y-axis represents the - Log10 p-value. Each point corresponds to a single microbial gene. The horizontal dashed line represents the nominal significance threshold (p = 0.05). No microbial genes were found to be significantly differentially expressed between T1 and T2 after correction for multiple comparisons (adjusted *p* < 0.05). c) Analysis of the composition and expression dynamics of the subgingival metatranscriptome using the custom database pipeline. The Venn diagram (left) illustrates the distribution of microbial genes detected at T1 and T2, highlighting a large core metatranscriptome of 20,687 genes (66.2%) shared between both timepoints. The bar chart (right) displays the top 20 microbial genes ranked by the magnitude of absolute Log2 Fold Change in expression between T2 and T1, as determined by DESeq2 from the custom database analysis. Genes with higher expression at T1 are shown in blue, while those with higher expression at T2 are in red. Consistent with the traditional pipeline differential expression analysis, none of these transcriptional changes were statistically significant (adjusted *p* > 0.05).

Next, we sought to identify specific microbial genes whose expression levels were significantly altered between baseline (T1) and the peak estradiol time point (T2). We employed a paired differential expression analysis using DESeq2, explicitly modeling the subject-specific effects. From the initial detection of over 3,108,995 gene features mapped to the HOMD reference database, a total of 172,851 genes passed quality and abundance filtering criteria and were included in the statistical testing. Despite the dramatic systemic and salivary hormonal changes observed, this analysis revealed no microbial genes that were significantly differentially expressed between T1 and T2 after correcting for multiple comparisons (Benjamini-Hochberg adjusted *p*-value < 0.05) (Figure 3B). Crucially, this lack of significant findings was replicated in the secondary pipeline analysis of the data mapped to the custom database, which also yielded no differentially expressed genes or taxonomic ranks (Supplementary file A). The absence of statistically significant differentially expressed genes (DEGs) across both pipelines suggests that the rapid, high amplitude increase in circulating estradiol, while detectable locally in saliva, did not elicit a strong, uniform transcriptional response at the individual gene level within the established subgingival microbial communities of this cohort.

To further characterize the extent of this functional stability, we examined the composition of the active metatranscriptome across both timepoints. A core of 20,687 microbial genes (66.2% of all detected features) was stably expressed at both T1 and T2, while 5,591 genes (17.9%) were detected exclusively at baseline and 4,955 (15.9%) exclusively at peak stimulation. The large shared core representing two thirds of the active metatranscriptome provides direct quantitative evidence of community-level transcriptional continuity across the hormonal perturbation. The genes detected exclusively at each timepoint showed no statistically significant differential expression and no enrichment in specific functional categories, consistent with stochastic detection variability rather than a directed biological response. Taken together, the absence of differentially expressed genes combined with the stability of the core metatranscriptome establishes functional resilience as a positive, measurable property of the established subgingival community under acute endocrine perturbation, rather than a null result.

## Discussion

Using controlled ovarian hyperstimulation as a defined experimental window, we demonstrate that the oral ecosystem exhibits a dual and dissociated response to acute supraphysiological estradiol: Biochemical reflection at the salivary level, functional resilience at the metatranscriptomic level, and selective ecological opportunism by *Prevotella* taxa. We leveraged the COH model, a unique human scenario involving rapid, supraphysiological E2 increases, to probe the resulting impact within the oral ecosystem. We focused on two distinct oral entities: saliva, as the immediate biochemical environment reflecting systemic hormonal influence, and the subgingival microbial community, to assess the functional transcriptional state of a key niche exposed to host vasculature. Our findings reveal a differential response within the oral ecosystem. While the systemic E2 increase was clearly mirrored in saliva, the subgingival microbiome exhibited a dual character: remarkable functional stability at the gene expression level, contrasted by a specific compositional response in key taxa sensitive to the hormonal surge.

Beyond the specific findings reported here, this study contributes a methodological framework with broader applicability. Controlled ovarian hyperstimulation provides a uniquely tractable human model for investigating acute hormone–microbiome interactions: the hormonal trajectory is pharmacologically defined, temporally bounded, clinically monitored with daily precision, and produces E2 surges that far exceed any naturally occurring physiological window. Unlike studies of chronic hormonal transitions such as menopause, contraceptive use, or hormone replacement therapy, COH compresses the endocrine perturbation into a two-week window with a known baseline, a quantified peak, and a defined endpoint, enabling genuinely causal temporal inference within a paired design. This experimental clarity is difficult to achieve in naturalistic hormonal research. The oral cavity, sampled here through two complementary matrices, saliva as a dynamic biochemical reflector and subgingival plaque as a stable biofilm niche, proved responsive at the hormonal level but resilient at the functional microbial level, a dissociation that would be impossible to detect without the temporal precision that COH affords. We therefore propose COH as a replicable human experimental platform for future mechanistic studies of hormone-microbiome crosstalk, applicable beyond oral health to any mucosal ecosystem accessible to paired longitudinal sampling.

The capacity of saliva to reflect systemic endocrine dynamics underpins its value as a non-invasive biomonitoring tool, particularly for steroid hormones like estradiol under physiological conditions (37). Our baseline findings empirically validate this principle, demonstrating a significant positive correlation between serum and salivary E2 levels within our cohort. This alignment with physiological expectations is robustly supported by literature spanning diverse hormonal contexts. Consistent serum-saliva E2 correlations have been reported not only during COH (12, 13, 38) but also across natural menstrual cycles and during hormone replacement therapy (39, 40). Fundamentally, this systemic-local linkage relies on the passive diffusion of the free, bioactive E2 fraction from the capillary beds surrounding salivary glands into the saliva (41). Therefore, our T1 data provide empirical support for salivary E2 as a reliable indicator of systemic bioactive hormone levels under physiological conditions.

On the contrary, the significant positive correlation between serum and salivary E2 observed at T1 diminished under the supraphysiological conditions of T2. This attenuation, where salivary E2 did not proportionally scale with markedly elevated serum E2, may stem from physiological adaptations during COH. For instance, high E2 levels during COH are known to increase hepatic sex hormone-binding globulin (SHBG) production (42). Elevated SHBG would reduce the free E2 fraction available for passive diffusion into saliva (43). Additionally, while passive diffusion across salivary glands is generally non-saturable (41), it is conceivable that extreme E2 concentrations could approach a transport or permeability threshold (44). Thus, the observed compression in salivary E2 at T2 likely reflects a combination of reduced free E2 bioavailability and potential limitations in trans-glandular passage at supraphysiological hormone levels.

Methodological factors related to E2 quantification likely also contributed to the observed attenuation in salivary E2 at T2. Our study utilized an enzyme immunoassay (ELISA) for salivary E2, a technique with a more limited dynamic range compared to mass spectrometry (LC-MS/MS). ELISAs can exhibit ceiling effects or signal compression at very high analyte concentrations, potentially underestimating true levels (45). Indeed, studies employing LC-MS/MS for salivary E2 measurement during COH have reported robust correlations at peak E2 levels, often without noting such pronounced compression effects (12, 13). Therefore, the choice of assay, particularly ELISA for samples with supraphysiological E2, may have influenced the apparent linearity of the serum-saliva E2 relationship at peak stimulation in our study.

Despite the complexities observed with absolute E2 values at T2, further analysis reinforces the utility of saliva for tracking COH-induced hormonal dynamics. Our robust linear regression, accounting for confounding factors such as gonadotropin dosage and stimulation duration, revealed a crucial insight. The magnitude of change in serum E2 significantly predicted the corresponding change in salivary E2. This finding strongly indicates that saliva effectively mirrors the intra-individual trajectory of E2 fluctuations. Indeed, even if the direct scaling of absolute E2 levels is compressed at peak stimulation due to physiological or technical limits, this result demonstrates that the pattern of change remains reliably reflected in saliva. This aligns with studies suggesting saliva’s utility for monitoring dynamic hormonal shifts (37). Thus, while caution is warranted when extrapolating absolute peak serum E2 from salivary measurements, saliva serves as a valuable tool for monitoring relative hormonal changes and individual response dynamics during ovarian stimulation.

In stark contrast to the measurable hormonal shifts, the subgingival microbiome exhibited notable functional stability between T1 and T2 despite the acute systemic and salivary E2 surge. Our analysis revealed no significantly differentially expressed genes, alongside highly personalized and temporally consistent transcriptional profiles for each participant. This observed resilience aligns with broader observations of the inherent stability of the oral microbiome in healthy individuals, even when facing host challenges (46–49). Specifically, the strong interindividual transcriptional signatures we noted, which overshadowed consistent time-related changes, resonate with studies highlighting personalized oral microbial ‘fingerprints’ (50–53), although investigations into metatranscriptomic responses to acute, high-amplitude hormonal shifts like COH remain limited. Therefore, our findings suggest that the established subgingival bacterial communities in this cohort were largely unperturbed at the gene expression level by the COH-induced hormonal changes within the investigated timeframe. Notably, this transcriptional stability extended to established virulence factors like microbial β-glucuronidases (54). This suggests that the acute E2 surge alone, in the absence of pre-existing periodontal disease, is insufficient to trigger the expression of microbial genes associated with chronic tissue degradation.

Several converging factors may explain this apparent lack of a gene-level transcriptional response within the subgingival microbiome. From a biological perspective, the established subgingival biofilm structure itself confers significant resilience against acute external perturbations (55). Additionally, the unique biochemical environment of the subgingival niche, bathed by serum-derived gingival crevicular fluid (GCF), may act as a buffer (56). This fluid could modulate the effective concentration or bioavailability of E2 reaching microbial cells, differing from systemic or salivary levels. Temporally, the short study window between T1 and T2, while capturing the hormonal peak, might simply be insufficient to trigger widespread, detectable transcriptional shifts within a mature, complex microbial community. Adaptive processes can require longer timescales or manifest more subtly. Finally, it also remains uncertain whether the bioactive E2 concentration within the GCF achieved a threshold high enough to trigger a robust transcriptional response across diverse microbial populations in this specific niche.

Beyond these biological and temporal considerations, inherent methodological and statistical aspects also contribute to the interpretation of these findings. The substantial interindividual variability in functional profiles, highlighted earlier, naturally makes it more challenging to detect consistent, potentially subtle treatment-induced effects across the cohort, even with a paired study design. Moreover, the modest sample size (*n*=10) limits statistical power, a critical factor in metatranscriptomic analyses that scrutinize tens of thousands of genes while demanding stringent control over false discoveries (FDR) (57). Consequently, it remains plausible that subtle or highly variable transcriptional responses occurred but did not meet the rigorous threshold for statistical significance within this study’s analytical framework.

A pivotal finding that nuances this narrative of functional resilience was the specific proliferative response of *Prevotella* species to the estradiol surge. This observation is mechanistically explained by the established capacity of taxa such as *Prevotella intermedia* to utilize host steroid hormones as a surrogate for their menadione (vitamin K) auxotrophy (58). Consequently, our results suggest that an acute E2 surge acts not as a community-wide stressor, but rather as a selective nutritional resource. This resource confers a distinct competitive advantage, fostering the growth of these estrogen-sensitive taxa. The direct correlation observed reinforces this as a clear localized adaptation to the endocrine microenvironment, positioning the oral cavity as a hormone-sensitive reservoir. The broader implication extends beyond oral health; such hormone-driven overgrowth may transiently increase the systemic load of microbial components like lipopolysaccharide (LPS) (59), representing a plausible pathway that contributes to the inflammatory etiology of distant, hormone-sensitive disorders such as polycystic ovary syndrome (60), endometriosis (61), and systemic autoimmune diseases (62) like rheumatoid arthritis.

Despite the insightful findings, this study has several limitations that should be considered. The sample size of ten participants restricts statistical power and the generalizability of our findings. The study duration between T1 and T2 captured only the acute hormonal response, potentially missing longer-term microbial adaptations. Our focus was solely on the subgingival niche; microbial communities in other distinct oral sites might respond differently to the hormonal stimulus. Metatranscriptomic analysis reflects RNA levels, which provide a snapshot of gene expression potential. However, these RNA levels may not perfectly correlate with downstream protein activity or ultimate community metabolic function. Direct E2 measurement within the GCF was not performed, leaving the precise local hormonal exposure within the subgingival niche unquantified. Finally, the use of ELISA for salivary E2 measurement introduces potential technical limitations regarding accurate quantification at the highest concentrations, as previously discussed.

### Conclusion

In conclusion, this study leverages the unique COH model to elucidate the response of the oral ecosystem to acute estradiol fluctuations. We demonstrate that salivary estradiol effectively captures the dynamics of systemic E2 surges, confirming its utility as a valuable tool for monitoring relative endocrine shifts within individuals. Concurrently, the established subgingival microbial community exhibits remarkable functional resilience at the transcriptional level, underscoring the inherent robustness of this niche. This highlights a dual response within the oral ecosystem: while the local biochemical environment sensitively reflects systemic hormonal changes, overall microbial community function maintains stability over acute timescales. However, this functional stability is nuanced by a specific compositional adaptation, where estrogen-sensitive *Prevotella* species leverage the hormonal surge for a competitive growth advantage.

Understanding this differential responsiveness is crucial for appreciating the complex interplay between systemic health and the oral environment. Future studies should expand this observational framework to additional mucosal sites, integrate GCF hormonal quantification, and increase cohort size to resolve the taxon-specific transcriptional responses that remain below the current detection threshold.

## Author contribution

Maria J. Rus: Conceptualization, Study design, Data acquisition, Formal analysis, Writing - Original Draft. Jack Lynch: Bioinformatic analysis, Formal analysis, Writing - Review & Editing. Ana T. Marcos: Patient recruitment, Data acquisition, Writing - Review & Editing. Thuy Do: Bioinformatic analysis, Writing - Review & Editing. David Alarcón-Alarcón: Formal analysis, Writing - Review & Editing. José Manuel Navarro-Pando: Patient recruitment, Clinical supervision, Writing - Review & Editing. Aurea Simon-Soro: Conceptualization, Study design, Supervision, Funding acquisition, Writing - Review & Editing. All authors gave their final approval and agreed to be accountable for all aspects of the work.

---

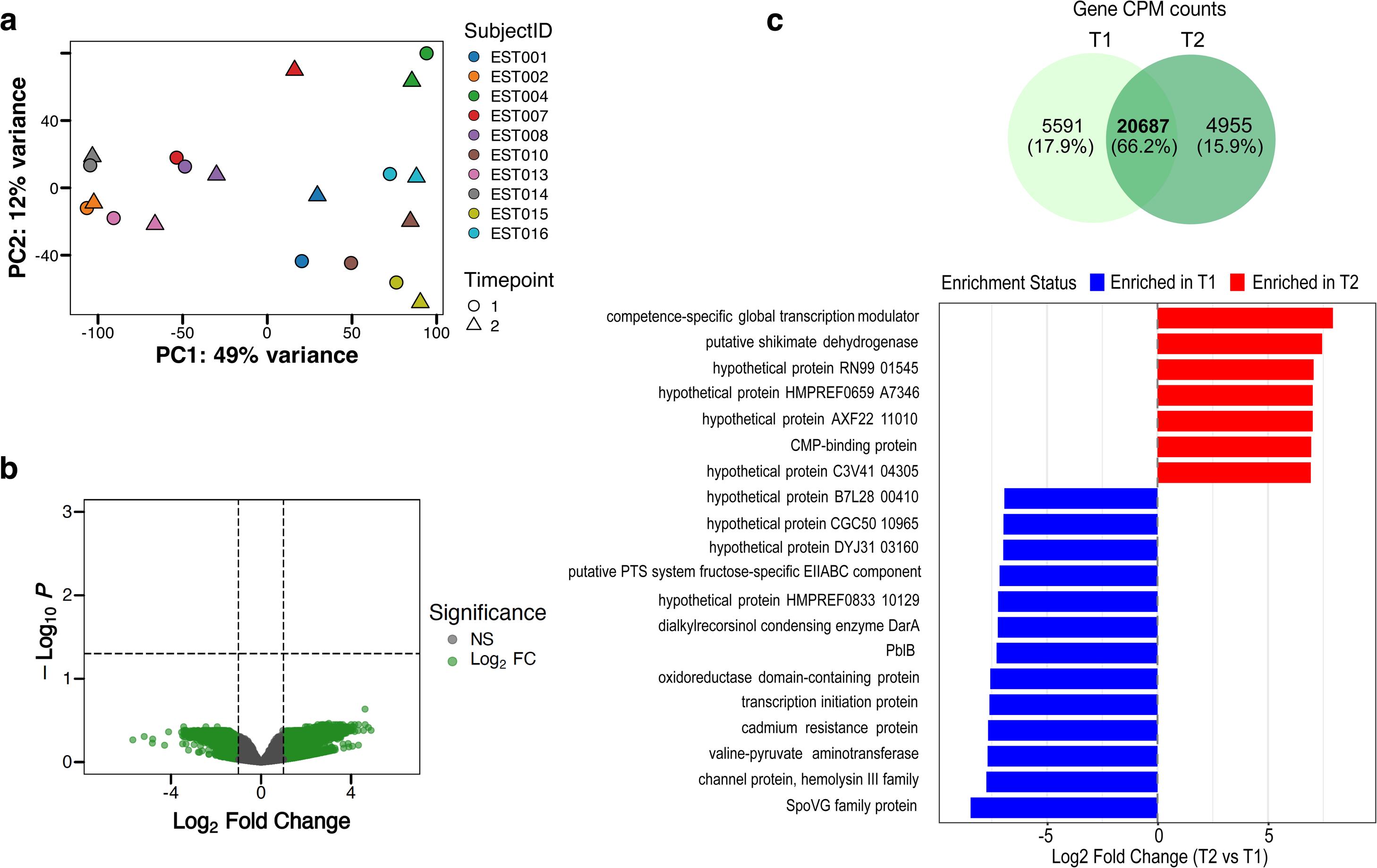

